# Evidence that polyploidy in esophageal adenocarcinoma originates from mitotic slippage caused by defective chromosome attachments

**DOI:** 10.1101/2020.05.14.096255

**Authors:** Stacey J. Scott, Xiaodun Li, Sriganesh Jammula, Ginny Devonshire, Catherine Lindon, Rebecca C. Fitzgerald, Pier Paolo D’Avino

**Affiliations:** Department of Pathology, University of Cambridge, Tennis Court Road, Cambridge, CB2 1QP, UK; Medical Research Council Cancer Unit, Hutchison/Medical Research Council Research Centre, University of Cambridge, Cambridge, UK; Cancer Research UK Cambridge Institute, University of Cambridge, Li Ka Shing Centre, Cambridge, UK; Department of Pharmacology, University of Cambridge, Tennis Court Road, Cambridge, CB2 1PD, UK

**Keywords:** cell division, chromosomal instability, kinetochore, genome doubling, PP1 phosphatase

## Abstract

Polyploidy is present in many cancer types and is increasingly recognized as an important factor in promoting chromosomal instability, genome evolution, and heterogeneity in cancer cells. However, the mechanisms that trigger polyploidy in cancer cells are largely unknown. In this study, we investigated the origin of polyploidy in esophageal adenocarcinoma (EAC), a highly heterogenous cancer, using a combination of genomics and cell biology approaches in EAC cell lines, organoids, and tumors. We found the EAC cells and organoids present specific mitotic defects consistent with problems in the attachment of chromosomes to the microtubules of the mitotic spindle. Time-lapse analyses confirmed that EAC cells have problems in congressing and aligning their chromosomes, which can ultimately culminate in mitotic slippage and polyploidy. Furthermore, whole-genome sequencing, RNA-seq, and quantitative immunofluorescence analyses revealed alterations in the copy number, expression, and cellular distribution of several proteins known to be involved in the mechanics and regulation of chromosome dynamics during mitosis. Together, these results provide evidence that an imbalance in the amount of proteins implicated in the attachment of chromosomes to spindle microtubules is the molecular mechanism underlying mitotic slippage in EAC. Our findings that the likely origin of polyploidy in EAC is mitotic failure caused by problems in chromosomal attachments not only improves our understanding of cancer evolution and diversification, but may also aid in the classification and treatment of EAC and possibly other highly heterogeneous cancers.

## Introduction

Genomic instability drives evolution, diversification, heterogeneity and adaptation in many cancers. One type of genomic instability, chromosomal instability (CIN), promotes large scale structural and numerical genomic changes that can lead to punctuated evolution by producing abrupt changes in the balance of critical growth and death pathways (1). CIN is usually classified as either structural - to indicate alterations in chromosome structure, such as translocations, inversions, duplications - or numerical - characterized instead by recurrent gain or loss of chromosomes that lead to aneuploidy and/or polyploidy. Some cancers present one specific type of CIN, but the two can occur together and numerical CIN can subsequently lead to structural chromosomal aberrations. Although both aneuploidy and polyploidy are very common in various cancers and numerical CIN is now considered a key factor in cancer evolution and diversification, its origin and exact role in cancer onset are still debated (1, 2). There is evidence that the presence of extra centrosomes, which is observed in various cancers, can directly lead to an increase in incorrect (merotelic) attachments of chromosomes to the spindle microtubules that in turn cause chromosome mis-segregation and aneuploidy (3). By contrast, our knowledge of the mechanisms that trigger polyploidy in cancer cells is still poor, despite the evidence that nearly 30% of different cancer types present whole genome doubling (WGD) events and that polyploidy propagates CIN, accelerates cancer genome evolution, increases tolerance to chromosome mis-segregation and drug treatments, and is associated with poor cancer prognosis (4–7).

Esophageal adenocarcinoma (EAC), the predominant histological type of esophageal carcinomas in the western world with high mutation burden and substantial heterogeneity (8–11), represents an ideal system to study the origins and role of polyploidy in cancer evolution and heterogeneity. EAC develops from a pre-cancerous condition known as Barrett’s esophagus (BE). BE can progress from a non-dysplastic lesion through intermediate stages of low-grade and high-grade dysplasia leading to EAC formation (12). During EAC development, the copy number and heterogeneity of the genome increases and the spectrum of mutations and rearrangements shows very little overlap with its paired BE counterpart (13). Whole genome sequencing (WGS) of paired BE and EAC samples indicated that, although approximately 80% of point mutations found in EAC samples are already present in the DNA from the adjacent BE epithelium (14), the difference in copy number aberrations between BE vs. EAC samples was much more dramatic. BE samples contained very few copy number changes and were mostly diploid, whereas EAC showed a wide range of copy numbers changes including some highly amplified regions (13, 15). These studies also found that up to two thirds of EACs emerged following a WGD event (tetraploidy), in a proposed pathway comprising an initial loss of p53 followed by tetraploidy and subsequent CIN (15).

There are four events that, in principle, can lead to polyploidy: cell fusion, genome endoreduplication, cytokinesis failure or mitotic slippage. In the latter case, cells fail to progress past metaphase and, after sustaining a prolonged arrest, the chromosomes decondense and cells re-enter in G1 phase. Both cytokinesis failure and mitotic slippage result in the formation of cells with a polyploid number of chromosomes, but the first leads to binucleate cells whereas the second to cells with large, tetraploid nuclei. As no clear evidence for either viral infection or endoreduplication has been reported in BE and EAC, we hypothesized that genome doubling in EAC development might arise as a result of a defect in cell division. To address this hypothesis and to understand the origin(s) of polyploidy in EAC, we analyzed cell division in both a 2D cell system that recapitulate EAC development and in patient-derived organoids. Our findings indicated that polyploidy in EAC originates from mitotic slippage caused by failure in chromosome alignment and segregation. Furthermore, WGS, RNA-seq, and quantitative immunofluorescence analyses suggested that an imbalance in the amount, and possibly regulation, of proteins involved in the attachment of chromosomes to spindle microtubules could be the molecular mechanism underlying this mitotic failure.

## Results

### p53-deficent BE and EAC cells display specific mitotic defects

As a first step to investigate the potential origin(s) of polyploidy in EAC, we analyzed cell division in a panel of BE and EAC cultured cells specifically selected in order to recapitulate the stages of progression from the pre-malignant condition to the carcinoma. We used two BE cell lines: CPA and CPD; the first is non-dysplastic, near-diploid, and has wild-type p53, while the second is dysplastic, near tetraploid and have mutated p53. We also analyzed four different EAC cell lines (all near-tetraploid and p53-deficient) and included the non-transformed immortalized RPE-1 cell line as a reference control cell line (Table S1). To analyze mitosis, cells were stained by immunofluorescence for the mitotic marker histone H3 pS10, tubulin and DNA, to calculate the frequency of mitotic cells or mitotic index (MI) and visualize the mitotic spindle and chromosomes. All BE and EAC cells had an MI lower than the control RPE1 cells (Fig. 1A). CPA, CPD and FLO cells had very similar MI values at around 4%, whereas JH-Eso-Ad1 and OE19 had slightly lower MIs, and OE33 cells had the highest MI among the esophageal cell lines, which was comparable to that of RPE1 cells (Fig. 1A). Quantification of cells at different mitotic stages (prophase/prometaphase, metaphase, anaphase and telophase/cytokinesis) revealed some interesting differences (Fig. 1B). As expected, in control RPE-1 cells the highest percentage of mitotic cells were found to be in prophase/prometaphase and telophase/cytokinesis (Fig. 1B). A similar trend was observed in the two BE cell lines and in OE19, albeit both CPA and CPD had a low percentage of cells in telophase/cytokinesis (Fig. 1B). By contrast, the other three EAC cell lines had a higher percentage of metaphase cells compared to RPE-1 and BE cells, which in FLO and JH-Eso-Ad1 was even higher than the number of cells in prophase. The combination in these EAC cells of an increase in metaphase cells without a high MI indicates that they spend long time in a metaphase-like configuration, possibly because of problems in aligning and/or segregating their chromosomes, but can then successfully progress through mitosis and the rest of the cell cycle. Immuno-fluorescence analysis also revealed that the p53-deficent CPD and EAC cell lines displayed a higher frequency of mitotic defects (up to 10%) than RPE-1 and CPA cells (Fig. 1C). A similar result was observed for the quantification of multinucleate cells (a readout for cytokinesis failure), although the frequency of this phenotype was much lower (<1.5%) (Fig. 1D). The mitotic defects could be categorized into three main phenotypes: lagging chromatin/chromosomes, multipolar spindles and failure in chromosome congression (Fig. 1E-G). Chromosome congression failure presented with varying levels of severity; in some cases the majority of the chromosomes successfully aligned at the metaphase plate with just a few chromosomes remaining at the spindle poles, whereas in other cells all chromosomes failed to congress and were randomly distributed over the mitotic spindle (Fig. 1F and G). We will hereafter collectively refer to chromosome congression defects as the ‘scattered chromosomes’ phenotype. This phenotype was the mostly frequent observed across all the p53-deficient cell lines and was absent in both RPE-1 and CPA. Multipolar spindles were also relatively frequent in FLO and OE33 cell lines. Lagging chromatin/chromosomes were observed in <1% of mitotic cells in all cell lines with the only exception of CPD.

**Fig. 1.**
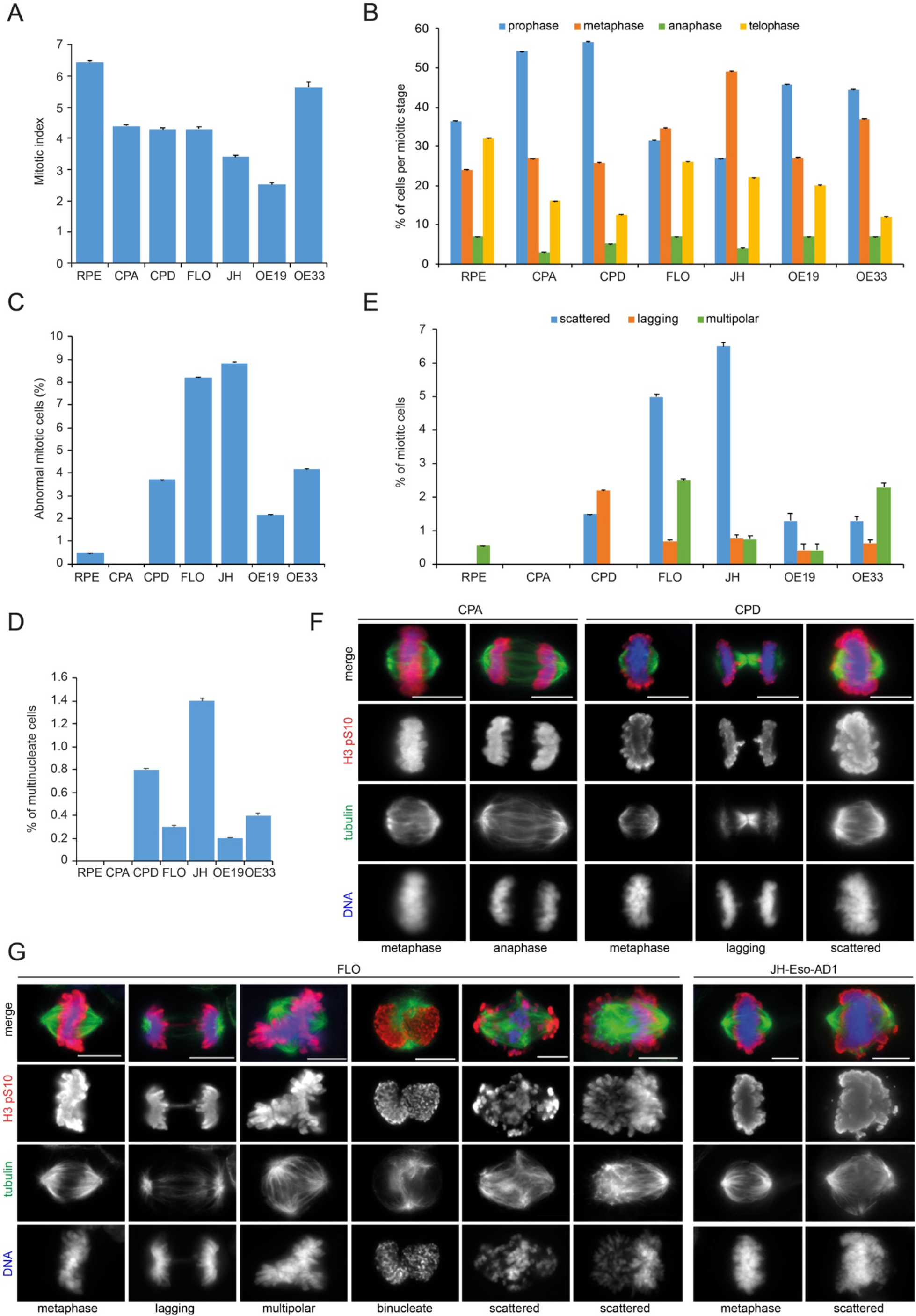
BE and EAC cells displays specific mitotic defects. (**A**) Cells from representative BE and OAC cell lines were stained to detect the mitotic marker histone H3 pS10, tubulin and DNA (see F and G below). The percentages of cells in mitosis (mitotic indices; MI) were counted (**A**) and categorized by each mitotic stage (**B**). In addition, for each cell line the number of abnormal mitoses was also counted (**C**) and categorized into one of three phenotypes: lagging chromatin, multipolar spindles or scattered chromosomes (**E**). (**D**) Graph showing the percentages of multinucleate cells from the experiments described in (**A-G**). More than 3000 cells in total and more than 200 mitotic cells per each cell line were counted; n≥10 independent coverslips. In each graph bars indicated standard errors. (**F-G**) Representative images from the indicated BE and OAC cell lines fixed and stained to detect the mitotic marker histone H3 pS10 (red in the merged images), tubulin (green in the merged images) and DNA (blue in the merged images). Bars, 10 μm.

A previous study reported that centrosome amplification occurred early in the progression of BE into EAC, and that this was dependent upon p53 loss (16). As supernumerary centrosomes can cause mitotic defects, we analyzed their presence in our BE cell lines and in the two EAC cell lines, FLO and JH-Eso-Ad1, that had the highest percentage of mitotic defects (Fig. 1E). We also asked whether extra centrosomes correlated with multipolar spindles and/or scattered chromosomes. BE and EAC cells were stained with antibodies against Plk4 and γ-tubulin to mark the centrioles and the respective centrosomes and then analyzed and quantified (Fig. S1). Both BE cell lines showed only bipolar spindles with two centrosomes and correctly aligned chromosomes, while FLO and JH-Eso-Ad1 cells had 10-11% of cells with more than two centrosomes (Fig. S1B), which often generated multipolar spindles, but with properly congressed chromosomes (Fig. S1A and C). Importantly, scattered chromosomes were only observed in cells with two centrosomes and bipolar spindles (Fig. S1C).

Together, our results indicate that p53-deficient BE and EAC cells have a significant increase in chromosome alignment defects that are not associated with extra centrosomes and multipolar spindles.

### EAC cells have a functional spindle assembly checkpoint and manifest mitotic slippage

We next employed time-lapse microscopy to better understand the origin of the mitotic defects in both BE and EAC cells and how they affected progression through mitosis. However, we first established whether these cell lines had a functional spindle assembly checkpoint (SAC). The SAC is a surveillance mechanism that monitors the attachment of chromosomes and prevent exit from mitosis until all chromatids have a correct bipolar attachment (17). In the presence of a functional SAC, cells arrest in mitosis when treated with the microtubule depolymerizing drug nocodazole. BE, EAC, and RPE-1 cells displayed variable increases in MI after nocodazole treatment, clearly indicating that they all possess a functional SAC (Fig. S2).

We incubated CPA, FLO and JH-Eso-Ad1 cells with the SiR-DNA dye to visualize chromosomes and then recorded images at five-minute intervals for 8 to 10-hour periods to monitor their progression through mitosis (Fig. 2). As expected, almost all CPA cells (90.0%; n=30) progressed through mitosis without problems (Fig. 2A, Supplementary Movie 1). A small percentage of CPA cells showed lagging chromosomes or failed cytokinesis (Fig. 2E), defects that were most likely caused by UV radiations or recording conditions. Similar to the results observed in fixed cells, 72% of FLO cells divided normally, while 6% formed multipolar spindles, 11% lagging chromosomes (Fig. 2B, Supplementary Movie 2), and 11% experienced mitotic slippage (Fig. 2E). In the last case, cells formed a broad metaphase plate, indicating chromosome congression defects, spent up to three hours in this configuration without entering anaphase, and then the chromosomes appeared to decondense and a nucleus reformed (Fig. 2C, Supplementary Movie 3). Mitotic slippage was also observed with even higher frequency in JH-Eso-Ad1 cells (28.6%; n=7) (Fig. 2D and E). Cells that failed to progress past metaphase appeared to have congressed their chromosomes at the equator, forming a well-defined metaphase plate, approximately 30 minutes after nuclear envelope breakdown (NEB), which marks the beginning of prometaphase. The cells then remained in this phase for about 90 minutes until the chromosomes began to drift away from the metaphase plate. In extreme cases, like the one showed in Fig. 2D, six hours after NEB the chromosomes drifted from the metaphase plate and attempted to realign, but failed and by the end of the recording period (8 hours) the cell begun to decondense the chromosomes, possibly because of cohesion fatigue (Fig. 2D, Supplementary Movie 4). The number of filmed JH-Eso-Ad1 cells was lower than the other two cell lines because most of the cells died during or prior to filming, despite our extensive troubleshooting to establish optimal conditions for these cells.

**Fig. 2.**
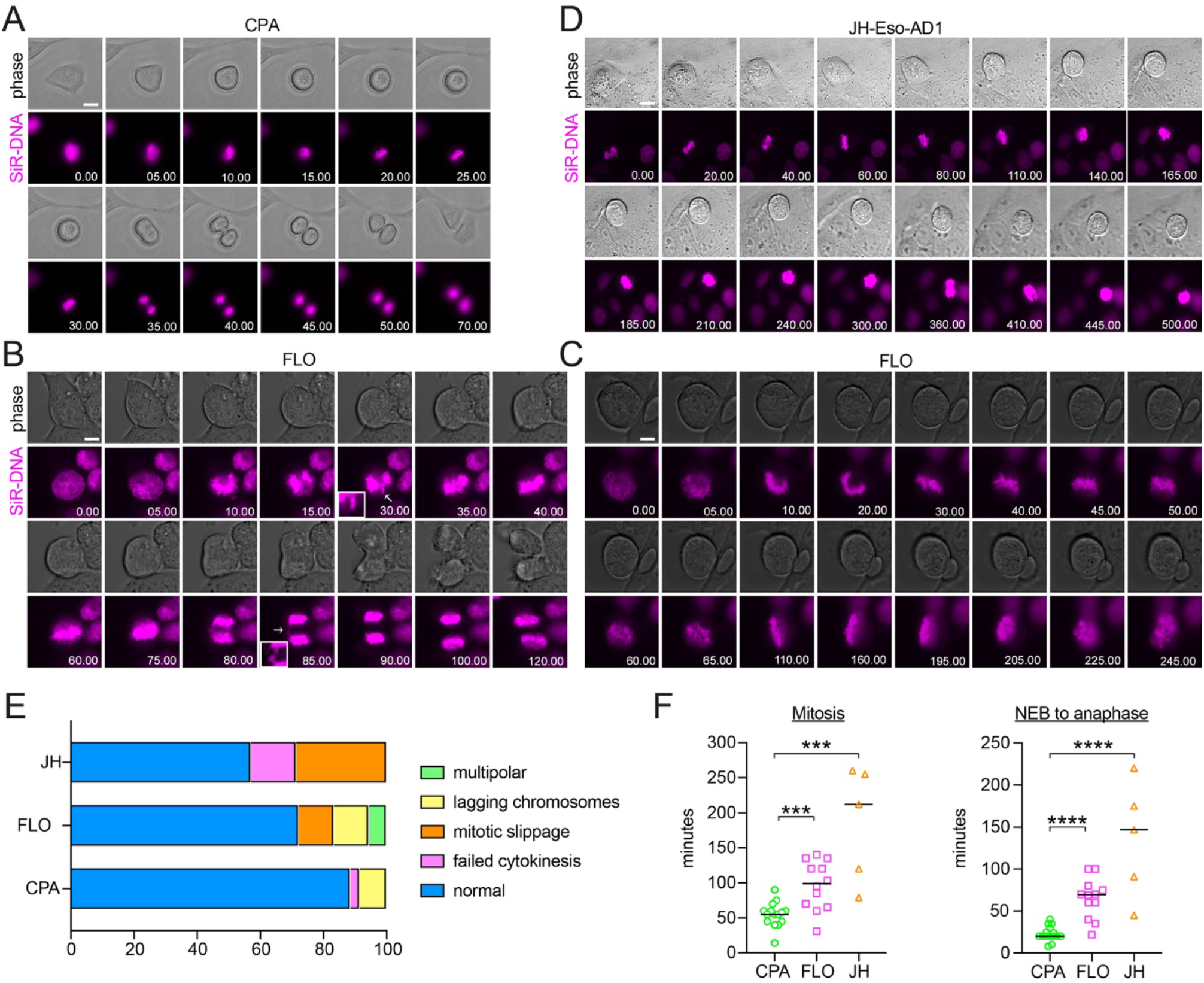
EAC cells have problems in congressing and aligning chromosomes and display mitotic slippage. (A-D) Images from time-lapse recordings of the indicated BE and EAC cells treated with SiR-DNA dye to visualize chromosome dynamics. Images were captured at 5 minutes intervals for 8-10 hours. Time is in min:sec relative to nuclear envelope breakdown (NEB). The arrows in (B) mark lagging chromatin. Bar, 10 μm. (E) Graph showing the frequency of phenotypes observed in the time-lapse recordings described in (A-D). 30 independent CPA cells, 18 independent FLO cells, and 7 independent JH-Eso-AD1 cells were analyzed. (F) Scatter plots showing quantification of the length of mitosis (from NEB until telophase) and the length of prometaphase-anaphase (from NEB until anaphase onset). n= 15 for CPA cells, n=12 for FLO cells, and n= 5 for JH-Eso-AD1 cells; *** *p* < 0.001, **** *p* < 0.0001 (Mann–Whitney *U* test).

**Fig. 3.**
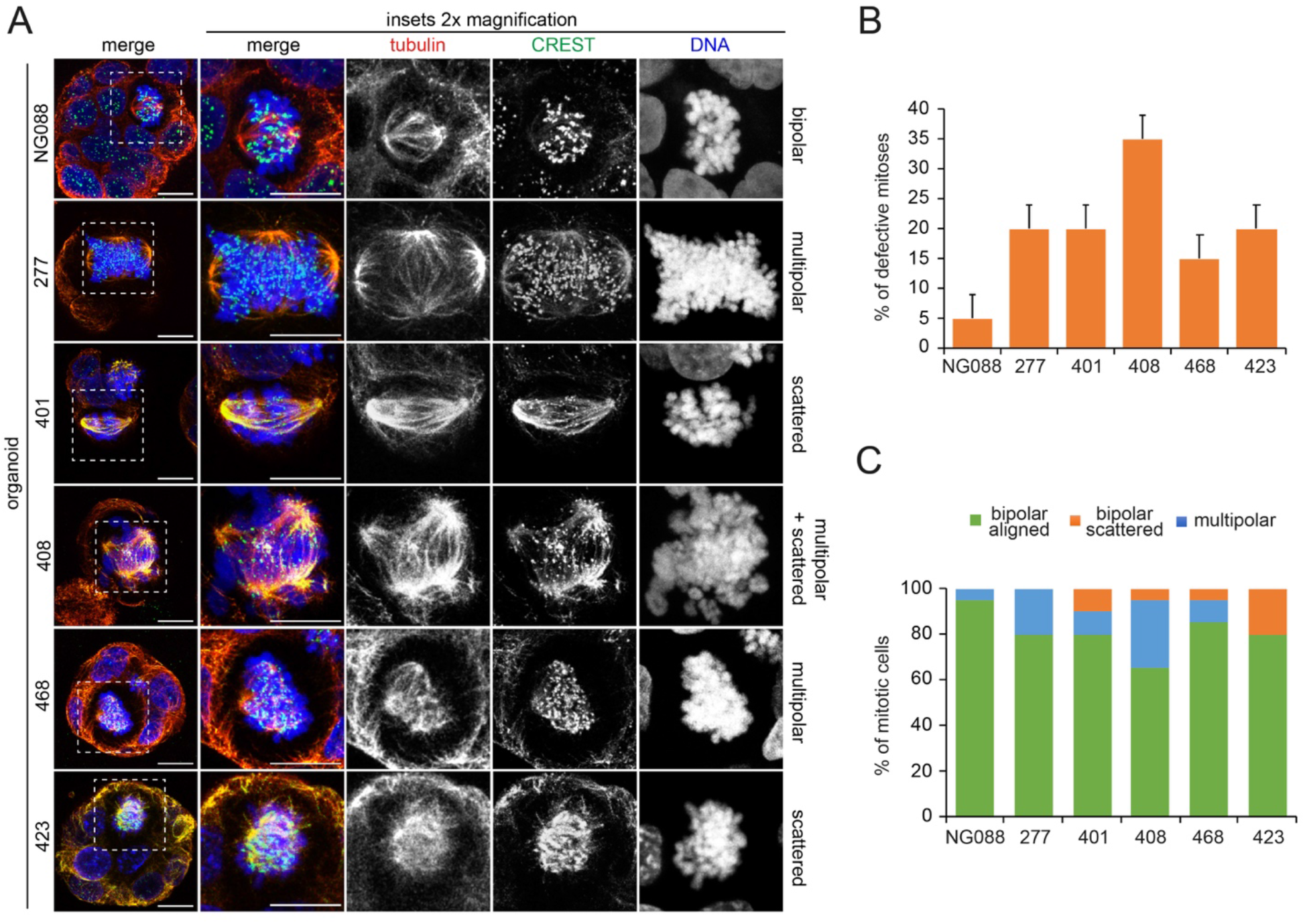
EAC organoids display mitotic defects similar to EAC cultured cells. (A) The indicated control and EAC organoids were fixed and stained to detect tubulin (red in the merged images), the centromeric marker CREST (green in the merged images), and DNA (blue in the merged images). Bars, 10 μm. (B) Graph showing the quantification of mitotic defects observed in the indicated organoids; n=20 independent organoids were analyzed for each group. (C) Graph showing the frequency of phenotypes observed in the organoids described in (A-B).

Our time-lapse experiments allowed us also to calculate the length of mitosis and we found that EAC cells took at least twice as long to divide than BE cells. Mitosis in CPA cells was completed in about 1 hour (58 ± 14 minutes), while it took an average of 103 (± 31) minutes in FLO cells and 212 (± 19) minutes in JH-Eso-Ad1 cells (Fig. 2F). It took also significantly longer for FLO and JH-Eso-Ad1 cells to reach anaphase onset after NEB than CPA cells (Fig. 2F), indicating that chromosome congression was more challenging in EAC cells.

In conclusion, these findings indicate that EAC cells have problems in congressing and aligning chromosomes, which can ultimately cause mitotic slippage and polyploidy.

### EAC organoids display mitotic defects similar to cultured cells

EAC cell lines are the most widely used model for EAC research, but in many cases we lack genomic information about both their primary tumors and germlines (18). Moreover, they can only recapitulate EAC to an extent because culturing them over numerous passages make them unrepresentative of the mutational features of the original tumor (18). To overcome these limitations, we have recently develop a series of EAC-derived organoids that more faithfully represent the primary cancers from which they derived and can be stably maintained (19). Therefore, we decided to further validate our findings from cultured cell lines by investigating whether EAC organoids presented similar mitotic defects.

A selection of organoids that represent healthy gastric tissues (NG088) and EACs with various ploidies and karyotypes (Table S2) were stained for tubulin, DNA and the centromeric marker CREST (20) to visualize, characterize and quantify mitotic figures (Fig. 4A). Almost all (95%, n=20) mitotic spindles observed in a normal gastric organoid line (NG088) were bipolar and had correctly congressed chromosomes (Fig. 4A and B). By contrast, all EAC organoids presented a much higher frequency of abnormal mitoses than NG088 (Fig. 4B). Similar to the data collected from cell lines, the most prevalent mitotic aberrations observed in EAC organoids were multipolar spindles and scattered chromosomes (Fig. 4C). With the exception of organoid CAM423, multipolar spindles were present in at least 10% of mitoses. The incidence of scattered chromosomes (5-20%) was sometime lower than multipolar spindles (Fig. 4C), but still comparable to the data obtained from EAC cell lines (Fig. 1). Interestingly, although these two mitotic defects were observed in both polyploid and aneuploid EAC organoids, one of the two polyploid organoids, CAM423, did not show multipolar spindles, while the other, CAM277, had no scattered chromosomes (Fig. 4C).

**Fig. 4.**
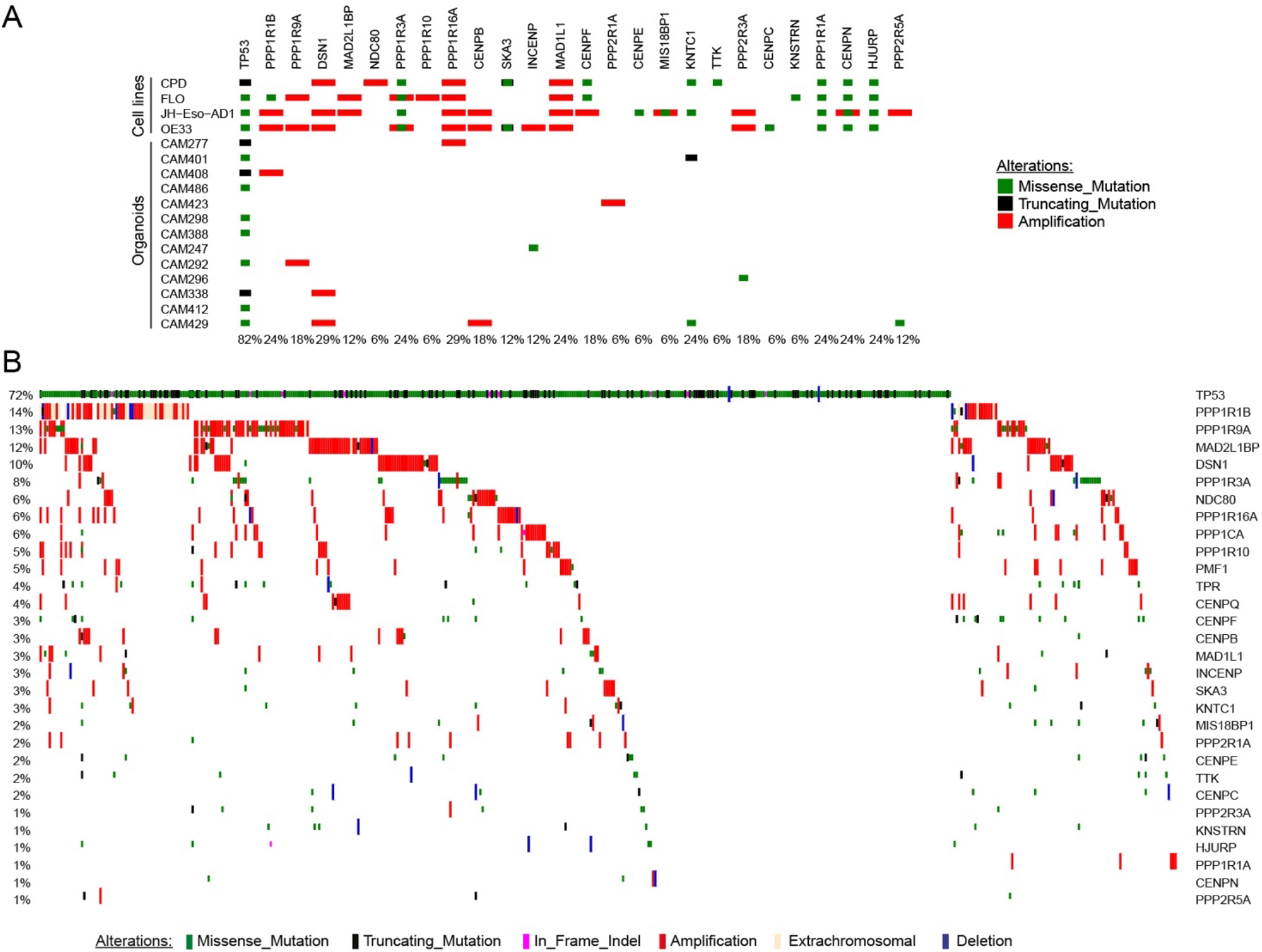
Genes involved in chromosome attachments are altered in EAC cells, organoids, and tumors. Diagrams showing the frequency and nature of SNVs, indels, and copy number alterations of genes involved in chromosome attachment in cell lines and organoids (A), and EAC cases (n=379) from the Oesophageal Cancer Clinical And Molecular Stratification (OCCAMS) consortium (B). Total proportion of cases altered by individual gene are listed at the bottom (A) or at the left (B) of the oncoplot.

Together, our findings indicate that EAC organoids present the same types of mitotic aberrations observed in EAC and BE cell lines. Although we could not find any clear correlation between mitotic defects and ploidy, the presence of scattered chromosomes did not appear to depend on the formation of multipolar spindles, which is consistent with the cell line data.

### Whole genome sequencing (WGS) of BE and EAC cell lines and primary tumors shows numerous copy number changes in kinetochore genes

The presence of chromosome alignment defects in both EAC cells and organoids suggested that the attachment of chromosome to microtubules might be impaired in these cells. The centromeric regions of chromosomes attach to microtubules through the kinetochore, a macromolecular structure composed of a multitude of proteins and protein complexes (21). The kinetochore is divided into two layers, the inner and outer kinetochore. The inner kinetochore comprises many CENP proteins that assemble onto the major centromeric protein CENP-A, a centromere-specific variant of histone H3, to form the constitutive centromere-associated network (CCAN) (21). The outer kinetochore is comprised primarily of the large multi-subunit Knl1/Mis12/Ndc80 complex network (KMN network), which is recruited by the CCAN at the inner kinetochore to form strong interactions with mitotic spindle microtubules (22). Furthermore, the association of kinetochores to microtubules is finely regulated by phosphorylation mediated by serine/threonine kinases and counteracting phosphatases, including Aurora B kinase (AURKB) (23).

We therefore investigated whether EAC cells, organoids, and tumors presented alterations in the structure and copy numbers of genes that might be responsible for the chromosome attachment defects observed in EAC cells and organoids. We selected genes encoding proteins known to be involved in chromosome alignment, including kinetochore components (e.g., CCAN and KMN proteins), SAC proteins, and regulatory factors, such as AURKB, Polo-like kinase 1 (Plk1), and members of the PP1 and PP2A phosphatase families (21, 24) (Table S3). WGS analyses revealed a number of alterations in our selected gene set that were shared across cell lines, organoids, and tumors (Fig. 4). The most common feature was gene amplification, which was frequently observed for the outer kinetochore components Dsn1 and Ndc80, the SAC proteins Mad1 and Mad2-binding protein MAD2L1BP (also known as p31 comet) (25), and various members of the PP1 family of phosphatases (Fig. 4). The picture for CCAN inner centromeric proteins (CENPs) was more mixed, with a combination of amplifications and missense mutations (Fig. 4). RNA-seq analyses confirmed that, in tumors, gene amplification corresponded to a significant increase in gene expression levels (p-value < 0.0001) for the mostly frequently amplified genes encoding for Dsn1, MAD2L1BP, and the regulatory PP1 subunit PPP1R1B (Fig. 5A). In many cases, PPP1R1B was over-amplified (ploidy adjusted copy number > 10), which is particularly interesting considering its close proximity to the oncogene ERBB2 that is also frequently over-amplified in EACs. Moreover, single sample gene set enrichment analysis showed a co-enrichment of our gene signatures with known cell-cycle hallmark gene signatures (Fig. 5B).

**Fig. 5.**
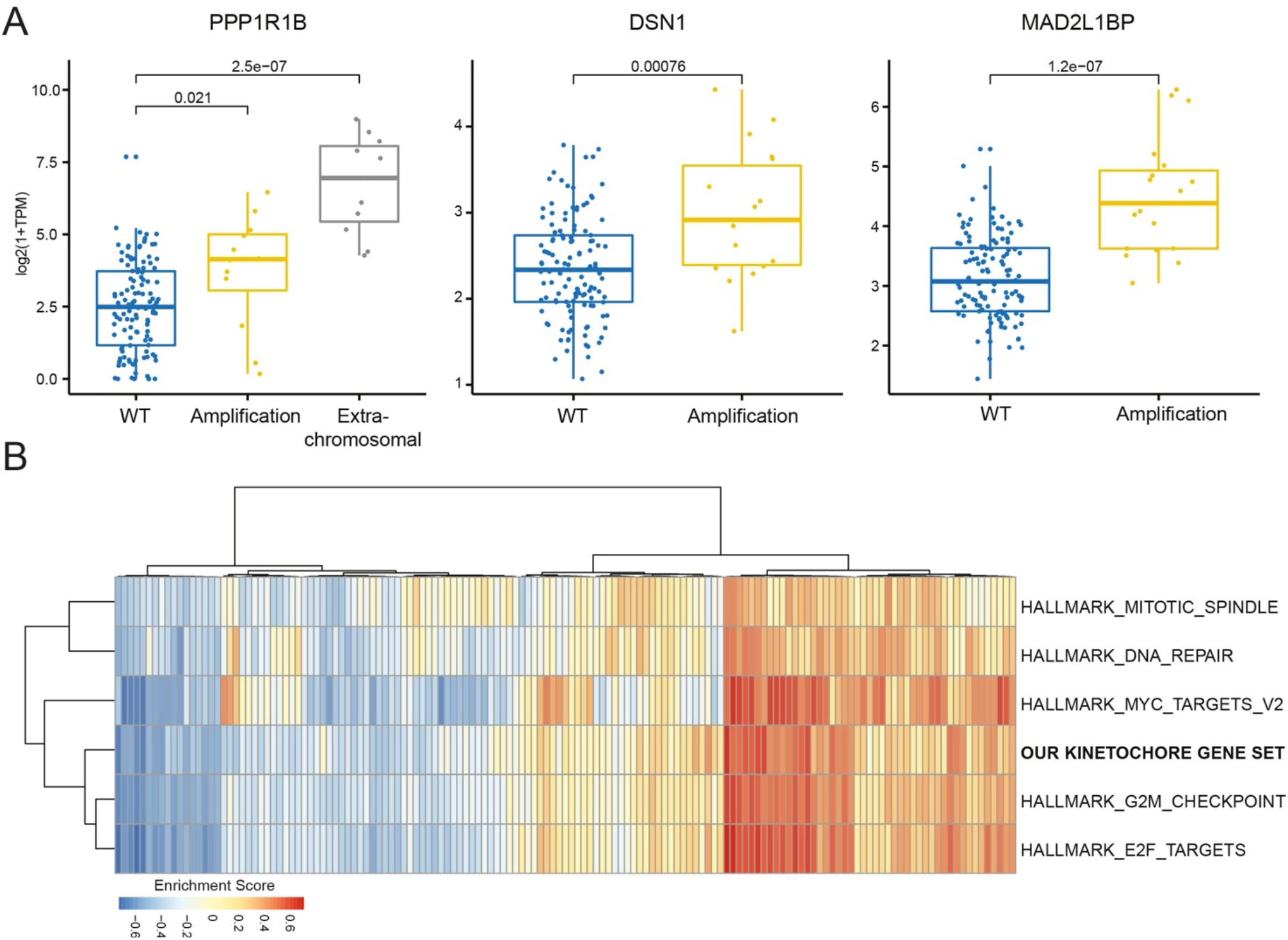
The expression of genes involved in chromosome attachments is altered in EAC tumors. (A) Graphs illustrating expression levels measured in log transformed Transcripts Per Million (TPM) for genes PPP1R1B, DSN1, and MAD2L1BP across EAC cases with different alteration types showing significant increase in expression upon amplification as compared to control wild type (WT) cases. (*p*-values computed through Wilcoxon test). (B) RNA-seq based single sample geneset enrichment analysis of annotated gene signatures across EAC cases (n=145) shows co-enrichment of our kinetochore gene signature with known cell-cycle hallmark gene signature (49).

These findings suggest that alterations in the copy number and expression of genes involved in chromosome attachment might be connected to the mitotic defects observed in both cell lines and organoids.

### The expression levels of inner and outer kinetochore components are altered in EAC cells

The alterations in copy number and expression of our gene set highlighted by the WGS and RNA-seq analyses prompted us to investigate whether these changes were also reflected at protein level in BE and EAC cells. We first analyzed the levels of a selection of SAC and kinetochore proteins, BubR1, CENP-C, Dsn1, Ndc80, and Spc24, by Western blot in unsynchronized BE and EAC cells (Fig. 6A). CPA cells were used as reference controls because are not dysplastic and have very few copy number changes (26) (Table S1). In agreement with our WGS and RNA-seq results, Western blot analysis revealed substantial differences in the levels of kinetochore proteins both within individual cell lines and comparatively across the different BE and EAC cell lines. The levels of the outer kinetochore components Ndc80 and DSN1 (member of the Mis12 complex) were increased between two- and four-fold in all cell lines compared to the CPA (Fig. 6A). The levels of the outer kinetochore component Spc24 (member of the Ndc80 complex) and inner kinetochore component CENP-C showed a much more modest increase and only in FLO and JH-Eso-Ad1 cells (Fig. 6A). By contrast, the mitotic checkpoint protein BubR1 showed slightly decreased expression in CPD, FLO, and JH-Eso-Ad1 cells. Interestingly, an additional, slower migrating BubR1 band was present in CPD and, more weakly, FLO cells, which could represent either a longer isoform or a specific post translational modification (Fig. 6A).

**Fig. 6.**
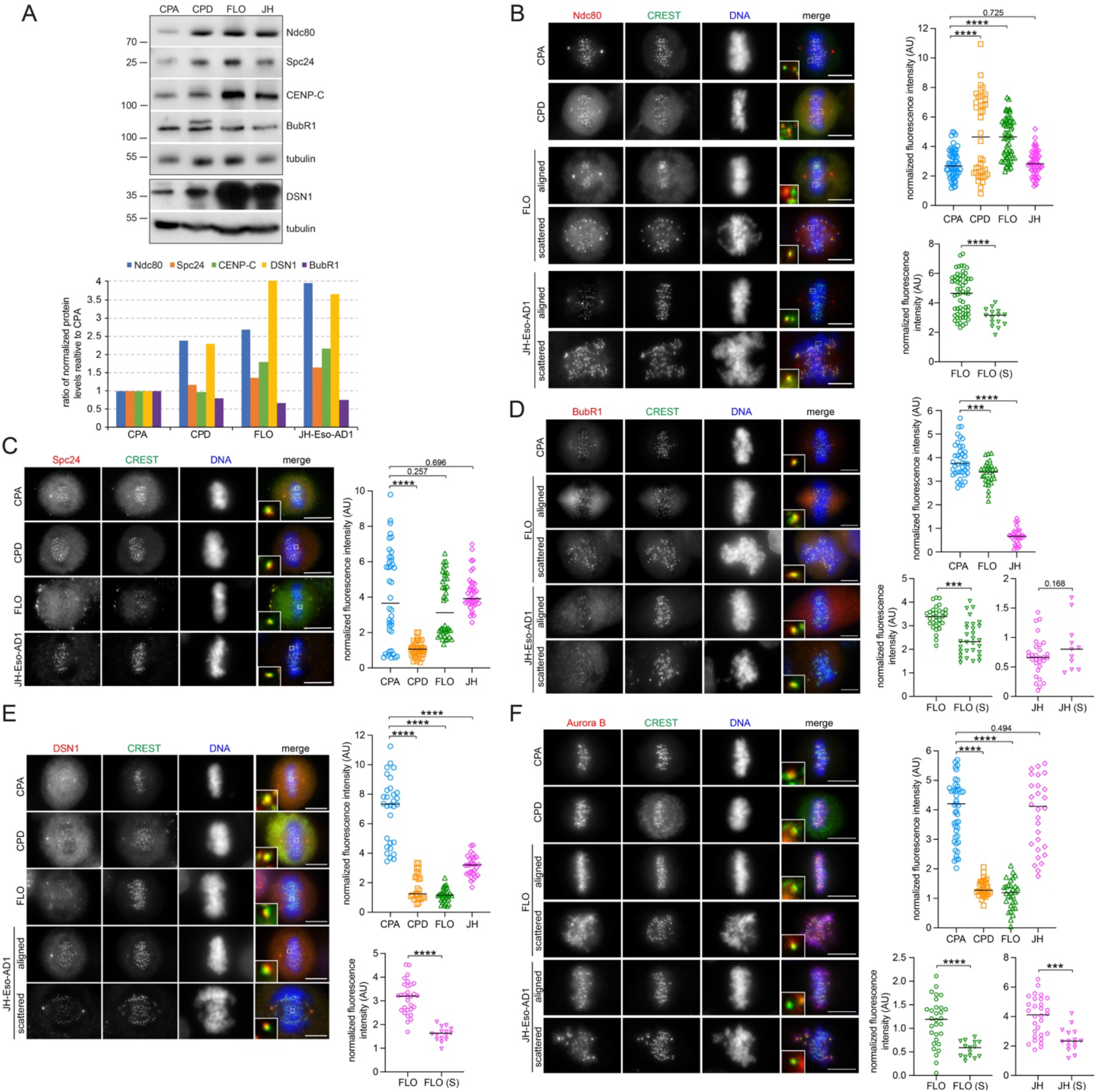
Kinetochore protein levels are altered in dysplastic BE and EAC cells. (A) Analysis of protein expression in BE (CPA and CPD) and EAC cells (FLO and JH-Eso-AD1). Proteins extracts from the indicated BE and EAC cells were separated by SDS PAGE and analyzed by Western blot to identify the proteins indicated to the right. The numbers on the left indicate the sizes of the molecular mass marker. The graph at the bottom shows the quantification of protein levels, normalized to tubulin and relative to levels in CPA cells. (B-F) Indicated BE and EAC cells were fixed and stained to detect the indicated epitopes. The insets show a 5x magnification of selected kinetochores. Bars, 10 μm. In each panel, the graphs to the right show quantification of fluorescence levels normalized to the centromere marker CREST (see Methods). More than 25 kinetochores from cells with aligned chromosomes and at least 10 kinetochores from cells with scattered (S) chromosomes were counted in n≥3 independent experiments. Horizontal bars indicate medians; **** *p* < 0.0001, *** *p* < 0.001 (two-tailed unpaired student’s T-test).

We next analyzed the accumulation of kinetochore proteins in mitotic cells by immunofluorescence. In every experiment, the level of each kinetochore protein was normalized to the level of a centromeric marker derived from human CREST patient serum (20) (see Methods). The accumulation of Ndc80, Spc24 and BubR1 at kinetochores followed in large part the same profiles observed in our Western blot analysis (Fig. 6B-D). The kinetochore intensity of Ndc80 in CPD and FLO cells was significantly higher than in CPA cells, although no significant difference was observed between JH-Eso-Ad1 and CPA controls (Fig. 6B). By contrast, the levels of the other Ndc80 complex subunit, Spc24, in EAC cells were not significantly different from CPA controls, while there was much less accumulation of Spc24 in CPD cells (Fig. 6C). In full accordance with the Western blot analysis, BubR1 levels were reduced in both EAC cell lines (Fig. 6D). Unfortunately, we failed to obtain a clear BubR1 signal in CPD cells, which might be possibly related to the presence of an additional band in Western blots (Fig. 6A). In striking contrast to the Western blot analysis, the levels of the Mis12 subunit DSN1 were significantly reduced in all cell lines compared to CPA controls (Fig. 6E). We then analyzed the levels of AURKB because of its key role in correcting improper chromosome attachments (23, 27). AURKB levels were strongly reduced in both CPD and FLO cells, but not in JH-Eso-Ad1 (Fig. 6F). Finally, comparison between mitoses with aligned or scattered chromosomes revealed that the levels of all these kinetochore proteins were significantly reduced on kinetochores of scattered chromosomes in almost all EAC cell lines, with the only exception of BubR1 in JH-Eso-Ad1 cells (Fig. 6B-F).

In conclusion, these results indicate that both inner and outer kinetochore proteins show different expression and kinetochore accumulation levels in dysplastic BE and EAC cells compared to CPA controls, which is likely to contribute to the defects in chromosome attachments observed in these cells.

### EAC cells with scattered chromosomes have a significant increase of lateral kinetochore-microtubule attachments

Kinetochores are initially captured laterally on the wall of microtubules before an end-on conversion occurs (28). It is only in this end-on conformation that the growth and shrinkage of microtubules can impart the pushing and pulling forces required for chromosome congression (29). A marker of this conversion is the presence of the astrin-SKAP complex which is recruited to mature end-on kinetochores (30), the astrin subunit of this complex being required for the correct alignment of chromosomes (31). To investigate whether the chromosome congression defects observed in EAC cells could result from a failure in converting lateral to end-on attachments, we stained with antibodies against astrin to visualize and quantify the different types of kinetochore-microtubule attachments (Fig. 7A). In both CPA and FLO cells that had successfully congressed to the metaphase plate, less than 10% of kinetochores were laterally attached (Fig. 6B), whereas there was a two-fold increase of lateral attachments in FLO cells with scattered chromosomes (Fig. 7B). This data suggests that a problem in converting from lateral to end-on attachments could be in part responsible for chromosome congression failure in EAC cells.

**Fig. 7.**
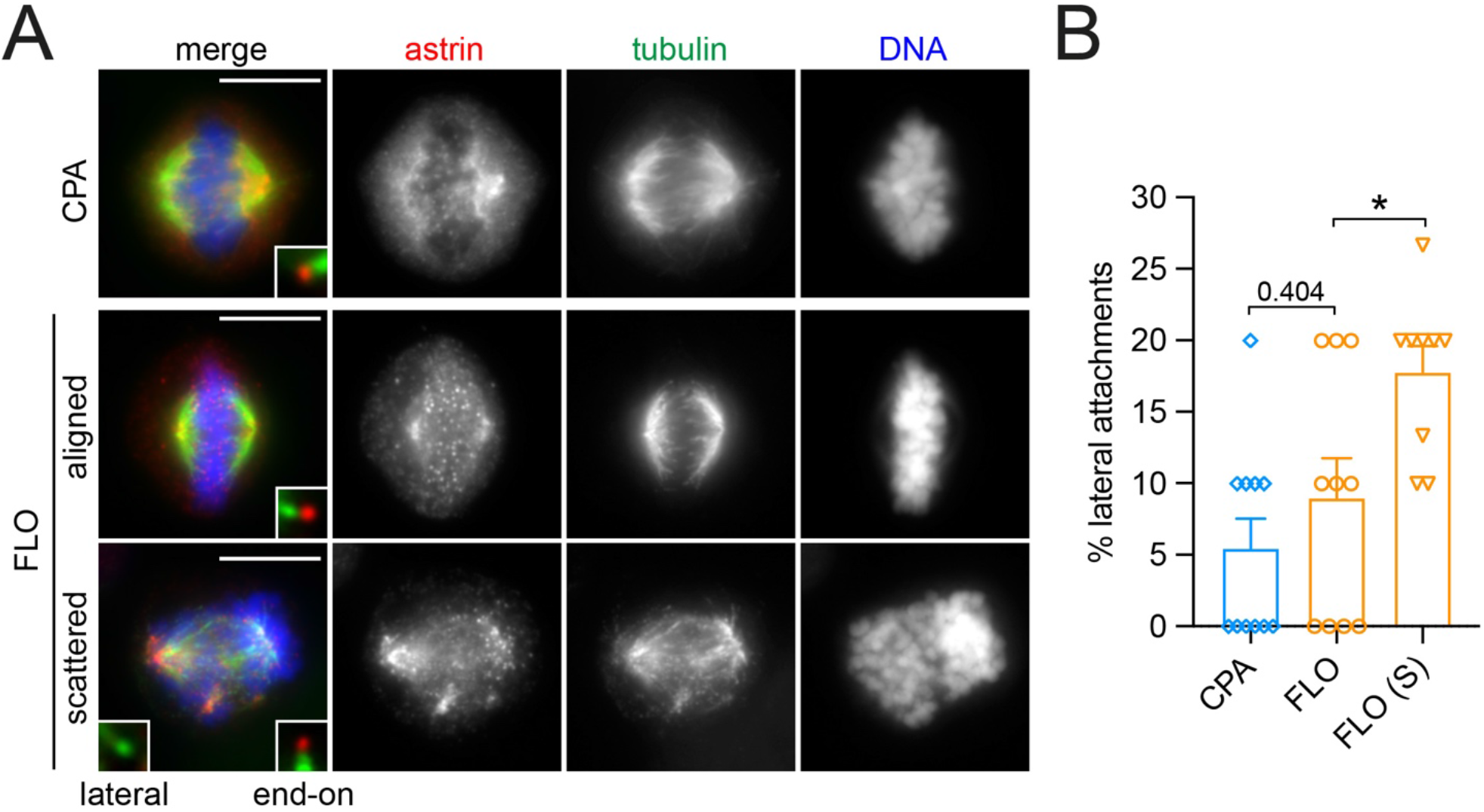
FLO cells with scattered chromosomes display an increase in lateral kinetochore-microtubule attachments. (A) CPA and FLO cells were fixed and stained to detect the indicated epitopes. The insets show a 5x magnification of selected kinetochores. Bars, 10 μm. (B) Graph showing the quantification of lateral kinetochore-microtubule attachments in CPA and FLO cells. 100 kinetochores were counted in each group; n=11 independent CPA cells, n=10 independent FLO cells and n=9 independent FLO cells with scattered chromosomes (S). Bars indicate standard errors; * *p* < 0.05 (Mann–Whitney *U* test).

## Discussion

The emerging evidence that polyploidy contributes to cancer evolution and heterogeneity by promoting CIN highlights the need to study the mechanisms that cause genome doubling in cancers not only to understand its role in cancer biology, but also to aid in the classification and design of therapeutic treatments of highly heterogeneous cancers. In this study we present evidence that polyploidy in EAC originates from mitotic slippage caused by defects in chromosome attachments during mitosis. Importantly, the frequency of these defects, less than 10% in cells and around 20% in organoids (Figs 1 and 3), is high enough to promote CIN, but not to significantly affect the viability of the entire cell population. We propose that these defects may permit the evolution of small clones within the large EAC cell population that can then more successfully adapt to the selective pressure of changing conditions.

The most prevalent mitotic defect observed in EAC cells was chromosome congression failure characterized by the presence of scattered chromosomes, which is a phenotype typically caused by defective kinetochores. Structural and/or functional alterations of kinetochore and centromeric proteins have been widely implicated in the promotion of chromosome mis-segregation and aneuploidy (32, 33). WGS and RNA-seq analyses coupled with analysis of kinetochore protein expression by Western blot and immunofluorescence revealed a number of copy number alterations and abnormal levels of numerous important constituents of the inner and outer kinetochore. Notably, our findings highlighted clear imbalances in the abundance of kinetochore proteins in BE and EAC cells. For example, both Western blot and immunofluorescence analyses indicated that the stoichiometry of the Ndc80 complex subunits Ndc80 and Spc24 was altered, as in most cases Ndc80 was more abundant than Spc24 in CPD and EAC cells (Fig. 6). Ndc80 overexpression has previously been described in brain, liver, breast, and lung cancers and increase in total Ndc80 translated into increased accumulation of this protein at the kinetochore in human colorectal carcinoma (HCT116), colorectal adenocarcinoma (HT29), osteosarcoma (U2OS) and cervix adenocarcinoma (HeLa) cells (34, 35). These findings led to speculate that increased accumulation of Ndc80, as part of the Ndc80 complex, might influence the interaction between kinetochores and microtubules in cancer cells. Our findings also support a similar conclusion that an increase of Ndc80 in EAC cells might be responsible for the chromosome alignment defects observed in these cells (Figs. 1 and 2). In addition, increase in Ndc80 was associated with a decrease in AURKB in both CPD and FLO cells (Fig. 6). The combination of reduced AURKB and Ndc80 overexpression could result in low Ndc80 phosphorylation and the formation of hyperstable kinetochore microtubule attachments and consequent problems in chromosome congression. Furthermore, the reduced levels of AURKB and BubR1 might weaken the SAC and allow mitotic slippage. It is important to point out that these alterations cause subtle changes in the regulation and mechanics of chromosome dynamics, which would allow most mitoses to progress normally. However, in a few cases these changes could lead to polyploidy and CIN, thereby promoting the evolution and diversification of new clones with potentially advantageous characteristics.

Our findings also indicated that the Mis12 component Dsn1 was amplified and expressed at high levels in both EAC cultured cells and tumors, but paradoxically its accumulation at kinetochore in mitosis was reduced (Figs. 4-6). We don’t have any explanation for this discrepancy at the moment, but these findings clearly indicate that a combination of different approaches - ranging from the analysis of genomes, transcriptomes, and proteomes down to the structure and composition of individual sub-cellular components - is necessary to fully evaluate the potential impact of gene and protein alterations in cancers.

One question raised from our data is whether the scattered chromosome phenotype occurs just because of chromosome congression failure or also because of cohesion fatigue after a prolonged mitotic arrest. The evidence that EAC cells take significantly longer than CPA to align their chromosomes (Fig. 2) would suggest problems in chromosome congression, although we cannot exclude that EAC cells take longer to align their chromosomes because they are polyploid. Moreover, our time-lapse experiments indicated that some EAC cells showed problems in maintaining chromosome alignment after mitotic arrest (Fig. 2 and Supplementary movies 3 and 4), which would suggest that the spread of the chromosomes over the mitotic spindle might also result from cohesion fatigue. Future time lapse experiments using multiple fluorescent markers, such as tubulin and cyclin B, could help resolve this issue.

Finally, our WGS analyses revealed that members of the PP1 family of phosphatases were frequently amplified in EAC cell lines, organoids, and tumors. In particular, PPP1R1B, also known as DARPP-32, was one of the genes that showed the highest increase in copy number in EAC patients. DARPP-32 is a neuronal protein and a potent PP1 inhibitor (36). Amplification of DARPP-32 and its truncated isoform t-DARPP has been found in 68% of gastric cancers and several studies indicated that it is also over-expressed in cancers of the breast, prostate, colon and esophagus - specifically in 30% of esophageal squamous cell carcinomas (37–43). PP1 phosphatases antagonize the phosphorylation of many mitotic proteins by AURKB (44) and it is therefore tempting to speculate that their de-regulation could affect kinetochore-microtubule attachments. Understanding how the PP1-AURKB regulatory balance is altered in EAC could provide extremely valuable insights for future targeted cancer therapies.

## Materials and methods

### Cell culture

RPE1 cells were obtained from ATCC and cultured in Dulbecco’s Modified Eagle Medium Nutrient Mixture F12 (Life technologies) supplemented with 10% FBS (Sigma) and 1% pen/strep (ThermoFisher). CPA and CPD cells were cultured in Keratinocyte-SFM supplemented with 2.5 μg prequalified human Epidermal Growth Factor 1-53 (EGF) (Life technologies), 25 mg Bovine Pituitary Extract (BPE) (Life technologies) and 0.5% penicillin/streptomycin (pen/strep) (ThermoFisher). FLO cells were cultured in Dulbecco’s Modified Eagle Medium (DMEM) (Sigma) supplemented with 10% Fetal Bovine Serum (FBS) (Sigma) and 1% pen/strep (ThermoFisher). JH-Eso-AD1 cells were cultured in Minimum Essential Medium (MEM) (Sigma) supplemented with 10% FBS and 1% pen/strep (ThermoFisher). OE33 and OE19 cells were cultured in Roswell Park Memorial Institute (RPMI) Medium (Life technologies) supplemented with 10% FBS (Sigma) and 1% pen/strep (ThermoFisher). All cells were cultured in a humidified atmosphere with 5% CO_2_ at 37°C.

### Organoid culture

Derivation and culture of organoids were as described (19). Briefly, organoid cells were resuspended in basement membrane matrix BME (Amsbio) and plated as 15 μl droplets. Once BME polymerized, culture medium was added and organoids were cultured in a humidified atmosphere with 5% CO_2_ at 37°C. Culture medium was refreshed every two days. To passage the organoids, BME (Amsbio) was disturbed by pipetting and organoids were disassociated using TrypLE (Invitrogen) at 37 °C.

#### Fluorescence microscopy

Cells were grown on microscope glass coverslips (Menzel-Gläser) and fixed in either PHEM buffer (60 mM PIPES, 25 mM HEPES pH 7, 10 mM EGTA, 4 mM MgCl2, 3.7% [v/v] formaldehyde) for 12 min at room temperature or in ice-cold methanol for 10 min at −20°C. For astrin staining, coverslips were first incubated for 5 minutes in pre-extraction buffer (60 mM PIPES pH 7.0, 25 mM HEPES pH 7.0, 10 mM EGTA, 4 mM MgSO_4_, 0.5 % [v/v] Triton X-100) and then fixed using PHEM buffer. After fixing, coverslips were washed three times for 10 min with PBS and incubated in blocking buffer (PBS, 0.5% [v/v] Triton X-100 and 5% [w/v] BSA) for 1 h at room temperature. Coverslips were incubated overnight at 4°C with the primary antibodies indicated in the figure legends, diluted in PBT (PBS, 0.1% [v/v] Triton X-100 and 1% [w/v] BSA). The day after, coverslips were washed twice for 5 min in PBT, incubated with secondary antibodies diluted in PBT for 2h at RT and then washed twice with PBT and once with PBS. Coverslips were mounted on SuperFrost Microscope Slides (VWR) using VECTASHIELD Mounting Medium containing DAPI (Vector Laboratories). Phenotypes were blindly scored by at least two people independently. Images were acquired using a Zeiss Axiovert epifluorescence microscope equipped with MetaMorph software.

Organoids were cultured on chamber slide for 7 days and washed twice with PBS. Cultures were fixed in 4% PFA at room temperature, quenched with 100 mM glycine-PBS for 10 mins, permeabilized in PBS + 0.5% Triton X-100 for 20 minutes, followed by blocking buffer (PBS, 1% BSA [w/v]) for 1 hour, and then incubated at 4°C with primary antibodies. After overnight incubation, organoids were washed and incubated with appropriate secondary antibody for 1 hour at room temperature as indicated above. Coverslips were mounted using VECTASHIELD Mounting Medium containing DAPI for nuclei staining. Organoids were imaged using a Leica confocal microscope TCS SP5, Z stacks were taken at 1-μm intervals and scored by two independent researchers. Images were processed by Volocity image analyse software (Perkin Elmer, version 6.3.0).

Fiji (45) was used to generate maximum intensity projections, which were adjusted for contrast and brightness and assembled using Photoshop. Fluorescence intensity values of kinetochore proteins were calculated using Fiji software and the following formula: (I_K_ - I_B_) - (I_C_ - I_B_) / (I_C_ - I_B_) = (I_K_ - I_C_) / (I_C_ - I_B_), where I_K_ = kinetochore intensity, I_C_ = CREST intensity, and I_B_ = background intensity.

### Time-lapse imaging

Cells were plated on an 8 well μ slide (Ibidi) in their appropriate growth media. Prior to recording media was replaced with Leibovitz’s L-15 media (ThermoFisher) containing 0.5 μM SiR-DNA dye (SpiroChrome). Cells were incubated in the dark at 37°C for 20 minutes and then imaged on an Olympus IX83 with an LED illuminator (Spectra-X, Lumencor), XY automated stage (ASI) in a 37°C incubation chamber (Digital Pixel) controlled using Micromanager freeware. Images were captured every 5 minutes for 100 frames with 100 ms brightfield and 20 ms Cy5 exposure times. Further processing was carried out using Fiji (45).

### Western blot

Cells were centrifuged, resuspened in phosphate buffer saline (PBS) and then an equal volume of 2x Laemmli buffer was added. Samples were then boiled for 10 min and stored at −20°C. Proteins were separated by SDS PAGE and then transferred onto PVDF membrane (Immobilon-P) at 15V for 1 hour. Membranes were blocked overnight at 4°C in PBS + 0.1% (v/v) Tween (PBST) with 5% (v/v) dry milk powder. After blocking, membranes were washed once with PBST and then incubated with the appropriate primary antibody diluted in PBST + 3% (v/v) BSA (Sigma) for 2 hours at RT. Membranes were washed 3×5 minutes in PBST and then incubated with HRPA-conjugated secondary antibodies in PBST + 1% BSA for 1 hour at room temperature. After further 3×5 min washes in PBST, the signals were detected using the ECL West Pico substrate (ThermoFisher) and chemiluminescent signals were acquired below saturation levels using a G:BOX Chemi XRQ (Syngene) and quantified using Fiji (45).

### Antibodies

The following antibodies and dilutions for Western blot (WB) and immuno-fluorescence (IF) were used in this study: mouse monoclonal anti α-tubulin (clone DM1A, Sigma, T9026; dilutions for WB 1:20000, for IF 1:2000), rabbit polyclonal anti-β-tubulin (Abcam, ab6046; dilutions for WB 1:5000, for IF 1:400), mouse monoclonal anti γ-tubulin (Sigma, GTU88; dilutions for IF 1:200), mouse monoclonal anti-cyclin B1 (Santa Cruz, clone GNS1, sc-245; dilution for WB 1:2000), mouse monoclonal anti-Aurora B (clone AIM-1, BD Transduction Laboratories, 611082; dilutions for WB 1:2000, for IF 1:100), rabbit polyclonal anti-phospho-histone H3 pS10 (Merck, 06-570; dilution for WB 1:10000 for IF 1:500), rabbit polyclonal anti-BubR1 (Abcam, ab172518; dilution for WB 1:10000, for IF 1:500), rabbit polyclonal anti-DSN1 (Sigma, SAB2702119; dilution for WB 1:2000, for IF 1:100), human anti-CREST (Antibodies Incorporated, 15-234; dilution for IF 1:1000) rat polyclonal anti-Plk4 (kind gift from P. Almeida Coelho and D. M. Glover, University of Cambridge, UK; dilution for IF 1:1000), rabbit monoclonal anti-Spc24 (Abcam, ab169786; dilution for WB 1:2000, for IF 1:200), mouse monoclonal anti-Ndc80/Hec1 (Santa Cruz, sc-515550, dilution for WB 1:500, for IF 1:50), rabbit polyclonal anti-astrin (Novus Biologicals, NB100-74638, dilution for IF 1:100). Peroxidase and Alexa-fluor conjugated secondary antibodies were purchased from Jackson Laboratories and ThermoFisher, respectively.

### Computational and statistical analyses

Mutation and copy number calls for different BE and EAC cell lines as published (18) were used for analysis. Copy number calling was performed by FREEC (46), as described (18). Mutations were called by GATK (Broad Institute, MA, USA), as described (18). Amplifications were defined as genes with 2× the median copy number of the host chromosome or greater.

To capture mutations and copy number alterations in EACs and primary organoids, we used WGS sequencing data of 379 EAC cases from the Oesophageal Cancer Clinical And Molecular Stratification (OCCAMS) consortium (8). We used final calls available for these datasets, where Strelka (47) was applied for calling SNVs/small indels and ASCAT (48) for copy number alterations. Genes were tagged as amplified if ploidy adjusted CN >2 and <10 or extrachromosomal-like if their ploidy adjusted CN >10. Along with WGS data, matched RNA-seq for subset of cases was used for transcriptomic analysis. We used processed expression quantification measured as Transcripts per Million (TPM) for analysis (8). Both genomic alterations and transcriptomic analysis was restricted to set of annotated kinetochore-related genes (Table S3). Only genes altered in at least three percent of cases were reported. To check whether amplification correlate with expression we compared expression of the target gene between cases without alterations and cases with amplification and level of significance was measured by non-parametric Wilcoxon test.

We ran single sample based gene set enrichment method to measure enrichment of annotated gene signatures (collection of genes). In this case we used known annotated hallmark gene signatures (49) related to cell cycle along with our annotated kinetochore genes and measured their enrichment across all EAC cases with RNA-seq data available using GSVA (50).

Unless otherwise specified, Prism8 (GraphPad) and Excel (Microsoft) were used for statistical analyses and to prepare graphs.

## Supporting information

Supplementary Figures S1 and S2, Supplementary Tables S1 and S2, Legends for Supplementary Movies S1 to S4

## Acknowledgments

We are very grateful to D.M. Glover for allowing us free access to his microscopy facility, and to M. R. Przewloka and P. Almeida Coelho for helpful suggestions and antibodies. We thank L. Capalbo for critical reading of the manuscript. SS was supported by an MRC PhD studentship. Work in PPD lab is supported by a BBSRC grant (BB/R001227/1). Work in CL lab is supported by BBSRC project grant BB/R004137/1. The laboratory of R.C.F. is funded by a Core Programme Grant from the Medical Research Council (RG84369). We thank the Human Research Tissue Bank, which is supported by the UK National Institute for Health Research (NIHR) Cambridge Biomedical Research Centre, from Addenbrooke’s Hospital. Whole genome sequencing was funded by a program grant from Cancer Research UK (RG81771/84119) to R.C.F.

