## Supplementary Figures S1 and S2, Supplementary Tables S1 and S2, Legends for Supplementary Movies S1 to S4 for "Evidence that polyploidy in esophageal adenocarcinoma originates from mitotic slippage caused by defective chromosome attachments"

**This PDF file includes:**

Supplementary Figures S1 and S2

Supplementary Tables S1 and S2

Legends for Supplementary Videos S1 to S4

**Other Supplementary Information for this manuscript include the following:**

Movies S1 to S4

Table S3

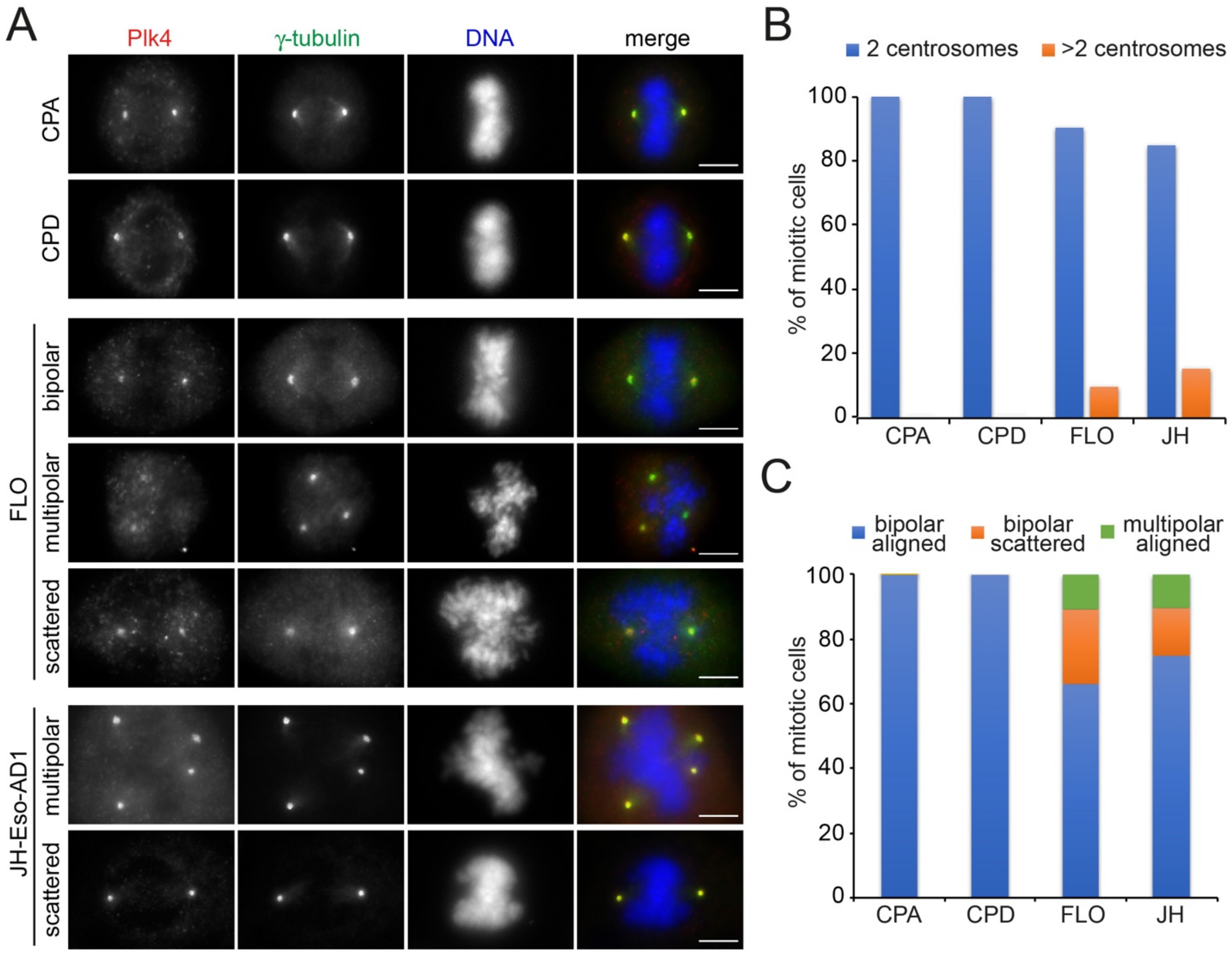


**Figure S1. EAC cells display extra centrosomes.** (**A**) Representative images from the indicated BE and EAC cell lines fixed and stained to detect Plk4 (red in the merged images), γ-tubulin (green in the merged images) and DNA (blue in the merged images). Bars, 10 μm. (**B-C**) Graphs showing the quantification of the number of centrosomes (B) and the defects (C) observed in in the mitotic cells from the experiments in (A). At least 50 mitotic figures from 3 different experiments were analyzed for each cell line.

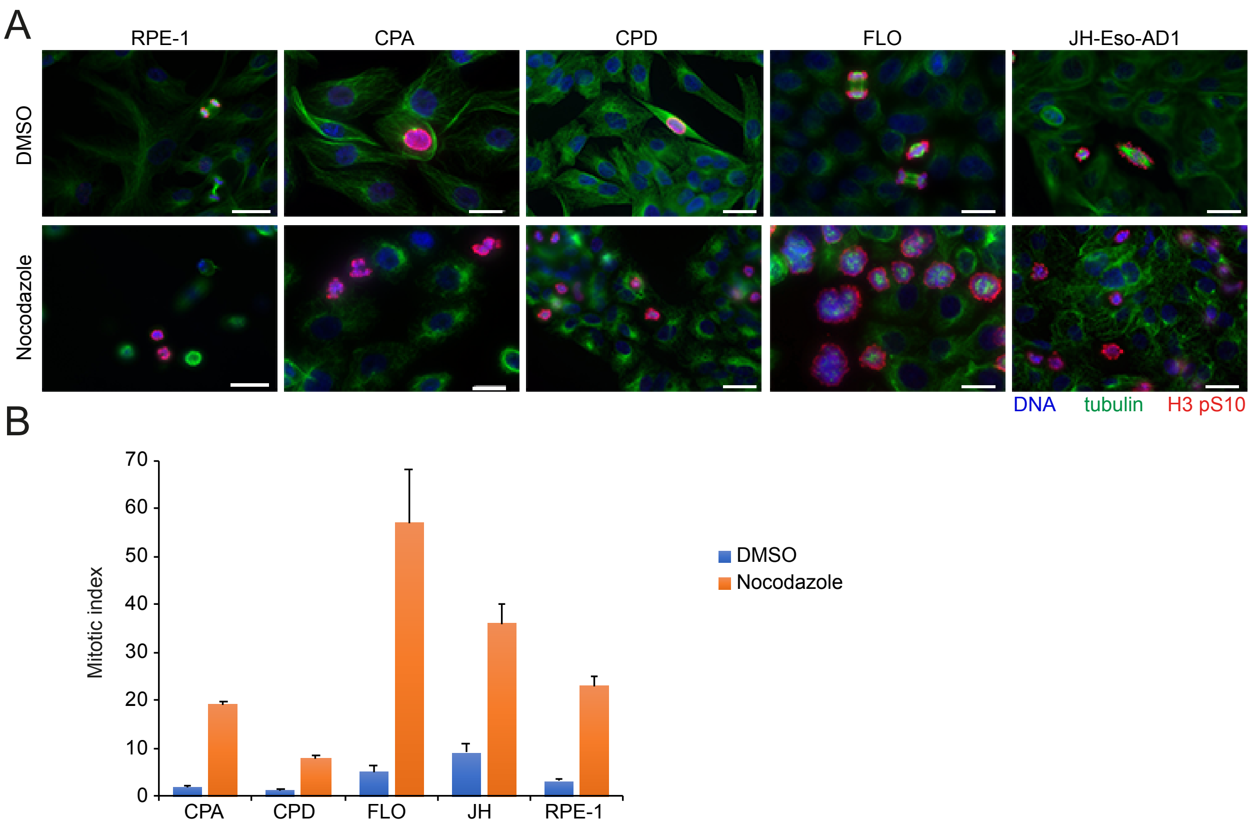


**Figure S2. BE and EAC cells have a functional spindle assembly checkpoint**. (**A**) Indicated BE and EAC cells were treated with the microtubule depolymerizing drug nocodazole or its solvent DMSO for 18 hours and then fixed and stained to detect the mitotic marker histone H3 pS10 (red in the merged images), tubulin (green in the merged images) and DNA (blue in the merged images). Bars, 10 μm. (**B**) Graph showing the quantification of cells in mitosis (mitotic indices; MI) for each cell line from the experiment described in (A). More than 1500 cells were counted for each cell line; n=6.

**Table S1. List of cell lines used in our experiments.**

| Cell line | Origin | Gender | Age | Ploidy | p53 | Reference /source |
|---|---|---|---|---|---|---|
| **RPE1** | Retinal epithelium, h-TERT immortalised | Female | Adult | Diploid | Wild type | (1)/ATCC |
| **CPA** | Non dysplastic BE, h-TERT immortalised | Male | Adult | Near diploid | Wild type | (2)/ATTC |
| **CPD** | Dysplastic BE, h-TERT immortalised | Male | Adult | Near tetraploid | c.404G>A | (2)/ATTC |
| **FLO** | EAC | Male | 68 | Near tetraploid | c.830G>T | (3)/EACC |
| **JH-Eso-AD1** | EAC stage 3, moderate to poor differentiation | Male | 66 | Near tetraploid | c.797G>A | (4)/to be deposited to ATCC |
| **OE19** | EAC stage 2, moderate differentiation | Male | 72 | Near tetraploid | c.929dup | (5)/EACC |
| **OE33** | EAC stage 2, poor differentiation | Female | 73 | Near tetraploid | c.404G>A | (5)/EACC |

**Table S2. List of organoids used in our analyses.**

| IDs | Age | Gender | Differentiation | p53 status | Ploidy |
|---|---|---|---|---|---|
| **NG088** | 76 | Male | Moderate | Wild type | Not known, but likely diploid |
| **CAM277** | 80 | Female | Poor | c.414delG, loss of p53 expression at protein level | Tetraploid genome, 100% of metaphases have a ploidy of >55. Ploidy of 4.04 indicating this organoid has experienced WGD. |
| **CAM401** | 77 | Female | Poor | Over-expressed p53 with hotspot mutation (R175H) | Aneuploid genome, ploidy of 1.74. Analysis of metaphases showed the following ploidy results: ~ 90% <40, ~5% 41-45 and ~5% 47-55. |
| **CAM408** | 60 | Male | Moderate | c.586G>A, loss of p53 expression at protein level | Aneuploid genome, analysis of metaphases showed the following ploidy results: ~10% <40, ~70% 41-45, ~5% 46 and ~5% >55. Overall ploidy score 1.94 |
| **CAM486** | 72 | Male | Moderate | Over-expressed p53 with mutation C.731C>T | Not determined |
| **CAM423** | 55 | Male | Moderate to poor | Wild type *TP53* and p53 expression pattern | Tetraploid genome. Ploidy 4.48 indicating this organoid has experienced WGD |

NG088 is the control organoid derived from non-cancerous cells from the stomach, the other organoids are all derived from EACs. All cultures are mixed heterogenous populations composed of cells with a variety of changes in chromosome number as well as chromosome rearrangements (6).

### Supplementary Movie Captions

**Supplementary Movie S1.**

This movie shows chromosome dynamics, visualized using SiR-DNA, in a CPA cell. All sequences were captured at 5 min intervals. Playback rate is 5 frames per second (FPS).

**Supplementary Movie S2.**

This movie shows chromosome dynamics, visualized using SiR-DNA, in a FLO cell. This cell completed mitosis, but showed lagging chromatin. All sequences were captured at 5 min intervals. Playback rate is 5 frames per second (FPS).

**Supplementary Movie S3.**

This movie shows chromosome dynamics, visualized using SiR-DNA, in a FLO cell. This cell failed to complete mitosis after an initial attempt to enter anaphase, and then chromosomes started to drift and decondense, indicating mitotic slippage. All sequences were captured at 5 min intervals. Playback rate is 5 frames per second (FPS).

**Supplementary Movie S4.**

This movie shows chromosome dynamics, visualized using SiR-DNA, in a JH-Eso-AD1 cell. This cell failed to complete mitosis after an initial attempt to enter anaphase, and then chromosomes started to drift and decondense, indicating mitotic slippage. All sequences were captured at 5 min intervals. Playback rate is 5 frames per second (FPS).
